# Shocker - a molecular dynamics protocol and tool for accelerating and analyzing the effects of osmotic shocks

**DOI:** 10.1101/2023.08.16.553535

**Authors:** Marco P. A. van Tilburg, Siewert J. Marrink, Melanie König, Fabian Grünewald

## Abstract

The process of osmosis, a fundamental phenomenon in life, drives water through a semi-permeable membrane in response to a solute concentration gradient across this membrane. In vitro, osmotic shocks are often used to drive shape changes in lipid vesicles, for instance, to study fission events in the context of artificial cells. While experimental techniques provide a macroscopic picture of large-scale membrane remodeling processes, molecular dynamics (MD) simulations are a powerful tool to study membrane deformations at the molecular level. However, simulating an osmotic shock is a time-consuming process due to the slow water diffusion across the membrane, making it practically impossible to examine its effects in classic MD simulations. In this paper, we present Shocker, a Python-based MD tool for simulating the effects of an osmotic shock by selecting and relocating water particles across a membrane over the course of several pumping cycles. Although this method is primarily aimed at efficiently simulating volume changes of vesicles it can handle membrane tubes and double bilayer systems as well. Additionally, Shocker is force field independent and compatible with both coarse-grained and all-atom systems. We demonstrate that our tool is applicable to simulate both hypertonic and hypotonic osmotic shocks for a range of vesicular and bilamellar setups, including complex multi-component systems containing membrane proteins or crowded internal solutions.

## Introduction

Osmosis is one of the most fundamental phenomena in life which occurs when a solute concentration gradient exists across a semi-permeable membrane, e.g., the lipid membrane of a biological cell. Due to the selective permeability of the membrane, solvent molecules diffuse along this osmotic gradient from a region of low solute concentration to high solute concentration, thereby equalizing the chemical potential in the two compartments. Osmosis is a vital process for a wide variety of organisms. In plant cells, for example, osmotic pressure leads to water influx into cells, which increases the internal pressure, so-called turgor pressure, against the cell wall. The turgor pressure controls the cell size, geometry, and rigidity which is required to maintain their cellular stiffness [1]. Bacteria, on the other hand, developed mechanisms to resist osmotic fluctuations, enabling them to survive in a wide range of environments and solute concentrations. By accumulating or synthesizing solute molecules, they maintain an internal solute concentration equivalent to the environment [2]. To cope with sudden solute concentration changes, bacteria use mechano-sensitive water channels such as MscL and MscS [3] acting as an osmotic pressure release valve.

Water movement across the plasma membrane of cells causes them to deflate or inflate, which is usually accompanied by shape changes. In vivo, membrane shape remodeling plays an active role in the regulation of many cellular processes including cell fission [4, 5]. Before cell fission can occur, an excess of membrane area is needed, which is obtained by lipid production and cell growth [ti]. The excess membrane is used to reshape the spherical cell into a dumbbell, a crucial intermediate for cell division events [7]. In experimental reconstitution studies, giant unilamellar vesicles (GUVs) are commonly used as (artificial) cell models [8-12]. The required excess membrane in these studies is obtained through decreasing the volume of the vesicle by a hypertonic osmotic shock, induced by a sudden change in the exterior solute concentration. To drive membrane fission, many cell and synthetic biology studies combined this approach of osmotic deflation with (i) encapsulation of scaffolding proteins or a protein machinery [13, 14] (ii) introduction of spontaneous curvature by protein crowding [15] or changes in pH [1ti]; (iii) insertion of lipophilic compounds [17, 18]; (iv) induction of liquid-liquid phase separation [19, 20]; (v) temperature cycling [21]; or (vi) microfluidic devices to apply a mechanical force [22].

Although the mentioned experimental studies led to promising fission results, the exact mechanism remains unknown since these fission events happen on a too short timescale to observe the process directly with sufficient detail. To overcome these shortcomings, computer simulations can be used to gain near to atomistic level resolution of membrane deformations at nano-to microsecond timescales [23]. Especially molecular dynamics (MD) simulations at the coarse-grained (CG) level have proven to be a valuable tool in studying vesicle shape transformations and fission [24-28]. Ghosh *et al*. [24], for example, constructed a range of spherical CG vesicles with different initial volumes and varying lipid ratios in the inner and outer leaflet to study differences in their shape deformations during volume reduction. They concluded that upon deflation, the initially spherical vesicles transformed into oblates, prolates, and dumbbells depending on the lipid ratio between the leaflets. In another approach, Markvoort *et al*. [2ti] simulated membrane fission pathways by distributing two non-miscible lipid types between the inner and outer leaflet starting from an already deformed vesicle. When both lipid types were distributed equally in the two leaflets, they observed pronounced phase separation. The resulting line tension caused a decrease in the neck radius of the initially dumbbell-shaped vesicle. Eventually, the authors observed asymmetric fission, i.e., the resulting daughter vesicles had a different lipid composition than the mother vesicle. However, when the two lipid types were distributed asymmetrically so that each leaflet contained only one type of lipids, an increase in spontaneous curvature led to symmetric fission.

The studies mentioned above show that MD simulations are suitable for examining the shape behavior of lipid vesicles during volume changes. However, to induce shape changes, i.e., change the initial area-to-volume ratio and generate an excess in membrane area, they rely on pre-deflated or pre-deformed vesicles instead of simulating an explicit osmotic shock. The reason is, that naively simulating an osmotic shock by letting solvent molecules diffuse along a concentration gradient through a membrane is too slow to induce any shape changes on typically accessible time scales in MD simulations. For example, Hong *et al*. [29] constructed a small atomistic bilayer consisting of 210 lipids and measured the water permeability of this membrane. After a 2 μs simulation 50 permeation events were counted, which is equivalent to 2000 permeation events in a vesicle with a radius of 1ti nm. Considering that such a vesicle contains around 150.000 water particles, in theory, a 75 μs simulation would be needed to yield a volume reduction of 50%. This calculation, however, does not account for any counter flux and the steady decrease in osmotic pressure upon water efflux. Therefore, realistic simulation times are likely in the range of several hundred microseconds.

While active water removal from the vesicle interior during the initial setup is a simple and suitable approach to avoid a time-consuming osmotic shock and yield general biophysical insights into their shape behavior, this approach allows very little control over obtaining specific shapes and deformation pathways. Considering the inherent slowness of osmotic shocks in vivo, removing a substantial amount of water instantaneously can alter the membrane dynamics resulting in non-physical shape deformation pathways. As demonstrated by Yuan *et al*. [30], when applying different volume-change rates to a mesoscopic model, different deformation pathways could be observed. For a slow rate, the membrane had sufficient time to relax the internal stress, resulting in prolate and dumbbell-shaped vesicles. On the other hand, a fast volume-change rate led to the accumulation of membrane stress, resulting in predominately biconcave oblate (discocyte) shaped vesicles. The use of pre-deformed vesicles as a starting point for further shape deformations could alleviate this problem, however, semng up realistic membrane shapes with a complex composition is currently still a non-trivial task [23]. Thus, a protocol for applying an explicit osmotic shock would offer so far unexplored opportunities, e.g., in studying the curvature sensing and remodeling behavior of proteins and lipids.

To accelerate osmosis in simulations and to allow beker control over the produced shapes, we developed a new MD protocol, which is implemented into the Shocker Python package. This protocol mimics the effects of water efflux (influx) during a hypertonic (hypotonic) shock by relocating solvent particles from the inner to the outer compartment (or vice versa). Although other protocols exist for removing or inserting solvent molecules [31, 32], there is, to our knowledge, no ready-to-go protocol for osmotic shocks, which is also compatible with the Martini CG force field. In addition to the protocol, we also implemented common analysis metrics to quantify vesicle shapes and shape changes such as the reduced volume and the reduced area difference which are explained in detail in the Supplementary Methods.

The remained of this paper is organized as follows: First, we describe the implementation of the solvent relocation scheme for mimicking osmotic shock exemplified by a hypertonic shock applied to a vesicle. Afterward, we consider four application systems: (1) hypertonic shocking of a simple vesicle, (2) hypotonic shocking of a crowded vesicle, (3) hypertonic shocking of a vesicle in the presence of proteins, and (4) hypertonic shocking of an atomistic double bilayers system. These systems serve to validate that the Shocker protocol can be applied to a wide range of biologically relevant systems and produces shape changes that are physically realistic and consistent with trends described in literature if available. Finally, we benchmark the performance and discuss limitations as well as future improvements.

### Algorithm

To simulate the continuous efflux (influx) of solvent molecules during an osmotic shock the simulation is divided into several pumping cycles. A pumping cycle includes the identification of solvent particles, relocating them into target bins, and updating the topology. In this section, we will provide a detailed description of the workflow of this cycle and discuss the underlying algorithms in six steps, which are visualized in **Figure 1**.

**Figure 1.**
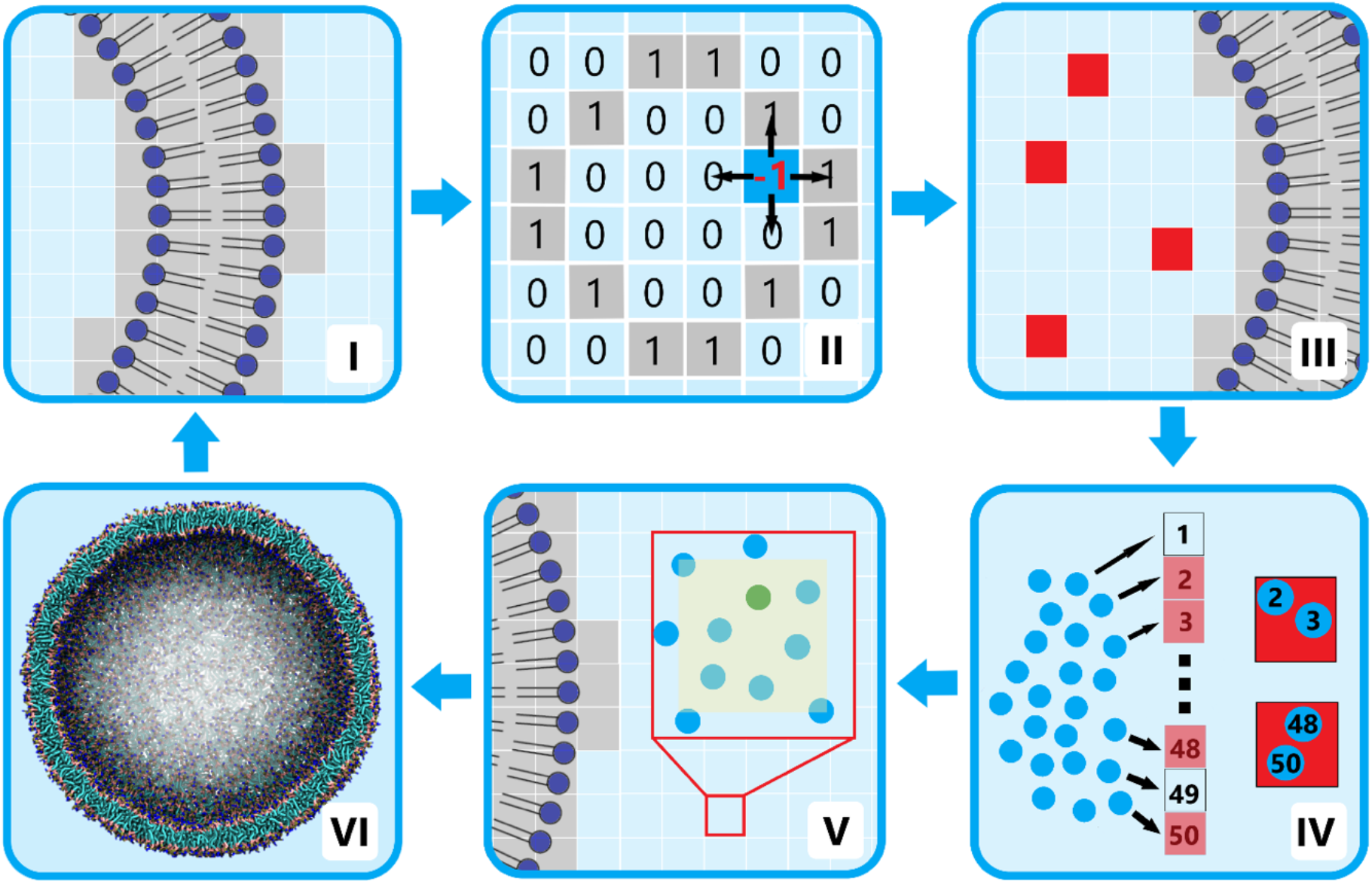
Workflow overview of one pumping cycle. (I) Initially, the original system is converted to a system of membrane bins (gray) and solvent bins (blue). (II) Subsequently, the solute bins are clustered using an algorithm based on graph clustering. (III) The solvent bins are randomly selected from the inner (outer) compartment for a hypertonic (hypotonic) shock, and (IV) the exact positions of solvent molecules in these bins are calculated. (V) One solvent molecule per bin is relocated to an ‘empty’ position in the other compartment after which a production run is initiated (VI).

In the following, a vesicle system is used as an example but the code is also applicable to other systems that feature an enclosed water compartment. In order to set up and initiate a pumping simulation the user should provide the structure file (GRO) of a fully energy-minimized and equilibrated vesicle. Besides, a topology file (TOP) and an index file (NDX) should be provided, where the index file should at least discern membrane and solvent particle groups. The online tutorials, deposited on our GitHub page (hkps://github.com/marrink-lab/shocker), describe more practical details concerning the input files and other requirements.

#### Step I: Binning the system

The pumping cycle starts with binning the system while conserving the system dimensions (**Figure 1**, step I), which drastically speeds up calculations compared to when considering each particle individually. The bin-edge size is adjustable, however, we recommend using the default bin-edge size of 1.3 nm, which works well for both AA and CG systems. To distinguish between membrane and solvent bins, the provided index file is used to translate the positions of all membrane particles to the bins they reside in, which receive the value 1. Subsequently, all remaining bins are set to 0 and are automatically part of the solvent group. Important to note here, is that all structures residing in the membrane such as proteins or cholesterol should be in the same index group as the lipids to avoid a non-contiguous bilayer bin system during the subsequent bin clustering process. The solvent group must solely contain the solvent particles, that are supposed to be relocated. Ions, soluble proteins, polymers, or any other molecule residing in the interior or exterior of the vesicle are omitted during the binning process.

#### Step II: Clustering of membrane and solvent bins

To discern the inner and the outer solvent compartment, the solvent bins are clustered using a distance-based clustering algorithm in which only adjacent bins are considered (**Figure 1**, step II) [33]. The algorithm starts in a bin with the value 0, indicating a solvent bin. From this position, it considers the values of the six adjacent bins, of which the position is added to the cluster if it turns out to be a zero. In **Figure 1**, step II these would be the bin below and left of the current bin. Subsequently, the current bin receives the value -1, indicating that this bin has already been visited, after which the algorithm proceeds to the next zero. If eventually, no adjacent zeros are available (the zero list is empty), the algorithm jumps to a new not processed zero, for as long as there are zeros in the box. In simple cases the user ends up with two clusters, however, in theory, there is a possibility to end up with more than two solvent clusters. This occurs in situations where the membrane crosses the periodic boundaries. In these cases, all smaller clusters not connected to the outer solvent cluster are assigned to the inner cluster. Consequently, Shocker only deals with assemblies that contain at most two water compartments.

The default bin-edge size is 1.3 nm. Working with a smaller bin size increases the precision as fewer solvent particles positioned close to the membrane surface are included in the membrane bin section. The downside of a smaller bin size is the increased probability of creating a non-contiguous membrane bin structure, ultimately causing the clustering algorithm to fail as all solvent molecules will form a single compartment. Using a smaller bin size also increases the total number of bins and therefore the overall computation time.

#### Step III: Solvent selection

The first step in solvent relocation is identifying those solvent particles that need to be relocated. Since the workflow example considers a hypertonic shock, the solvent is moved from the inner to the outer compartment. Bins are randomly selected from the clustered inner solvent compartment (**Figure 1**, step III). If a selected bin contains other molecules than just the solvent, it will be discarded and another bin will be selected. This ensures that the hydration shell of for example soluble proteins and polymers remains untouched. Only if no solvent-only bins can be identified, those bins will be considered further. From each of the selected bins, only one solvent molecule is chosen to be relocated to ensure that the solvent is removed from areas well dispersed across the whole inner compartment. The number of solvent molecules to be relocated (and hence the number of bins considered) is specified by the user.

#### Step IV: Particle identification

To identify the particles residing in the selected bins, the bin addresses of all solvent particles are compared to the addresses of the selected bins to retrieve the indices of the particles in these bins relative to the solvent list (**Figure 1**, step IV). Finally, these indices are converted to the global system indices to yield a list of indices per selected bin.

#### Step V: Solvent relocation

After selecting solvent particles, new positions have to be found in the target bins. Relocating particles by simply choosing random coordinates usually causes overlap between particles leading to numerical instabilities. Shocker’s relocating algorithm always tries to find the most suitable locations by taking the presence of other structures and particles into account (**Figure 1**, step V), thereby avoiding significant overlap.

The water placement starts by selecting a suitable bin for relocation, which ideally contains only solvent molecules. In case such bins are not available, a bin containing the smallest number of solute particles is selected. Subsequently, within this bin, a spot is identified with the largest distance to the nearest neighboring particles by performing trial insertions. To avoid overlap with particles residing in neighboring bins, this position has to be within 0.3 nm from the edges of the selected bin.

#### Step VI: Production MD run and steady-state

Once the desired number of solvent particles has been relocated a production run is initiated, which completes the pumping cycle (**Figure 1**, step VI). The length of these production runs is defined by the user and greatly depends on the type of system. For example, for a vesicle with a radius of 15 nm, we used a production run of 2 ns between each pumping cycle.

Besides hypotonic and hypertonic osmotic shocks, Shocker can perform a steady-state run, in which the volume of a vesicle, regardless of the shape, is kept constant. To this end, the number of enclosed water particles at the start of the steady-state run is calculated and compared to the current number after each cycle, which typically lasts 2 ns. Considering that a significant fraction of solvent molecules can reside in the interior of the membrane, they might belong to the vesicle interior in one cycle and to the exterior in the next one due to the rigorous clustering procedure in Step II. This means that the target number of solvent molecules can never be precisely determined. Consequently, translocated the exact difference in solvent molecules could lead to an overcompensation and thus further membrane deformation instead of a steady-state simulation. Therefore, the interior volume is averaged over 10 cycles, which means the volume is only adjusted after 20 ns.

Usually, the osmotic shock simulation runs until the desired number of pumping cycles is reached. The only cases in which the solvent pumping is completely terminated are when no solvent molecules are available for pumping or the membrane ruptures. However, the user can also define a target interior volume or solute concentration. Once this target is reached, Shocker switches into steady-state mode, and further pumping cycles are only initialized when the interior volume changes. To this end, Shocker monitors the solute concentration and interior volume if these options are switched on by the user. A detailed description of how these properties are obtained is given in the Supplementary Methods.

## Results

In the following, we apply an osmotic shock to five different example systems to validate Shocker’s algorithms. First, we simulated a simple CG vesicle composed of POPC lipids using the Martini 3 forcefield [34]. We compare the effect of different pumping rates on the deformation pathways during a hypertonic osmotic shock. Second, we performed a hypotonic osmotic shock on a large POPC vesicle until the vesicle ruptures. Third, we simulated a more complex CG vesicle system composed of a phase-separating lipid mixture (DPPC, DFPC, and cholesterol) that also contained a phase-separated polymer mixture (dextran and PEG) in the vesicle interior. Fourth, we studied a large CG POPC vesicle including a high concentration of aquaporin, a transmembrane protein. Lastly, to show Shocker’s compatibility with AA forcefields, we also performed a pumping simulation on a double bilayer POPC system using the CHARMM3ti forcefield [35]. More details about the simulation setups are provided in the Methods section and Supplementary Table 1.

### The pumping rate influences vesicle shape changes

In the first test case, we use Shocker to perform a hypertonic shock on a simple POPC vesicle with a radius of 7.5 nm filled with water molecules (**Figure 2b**). We examine the effect of different water efflux rates, which has been shown previously to result in different shape remodeling pathways [30]. Here, we used pumping rates of 0.1% (slow) and 1% (fast) of the initial vesicle volume, corresponding to the translocation of 10 and 100 water molecules per 2 ns, respectively.

**Figure 2.**
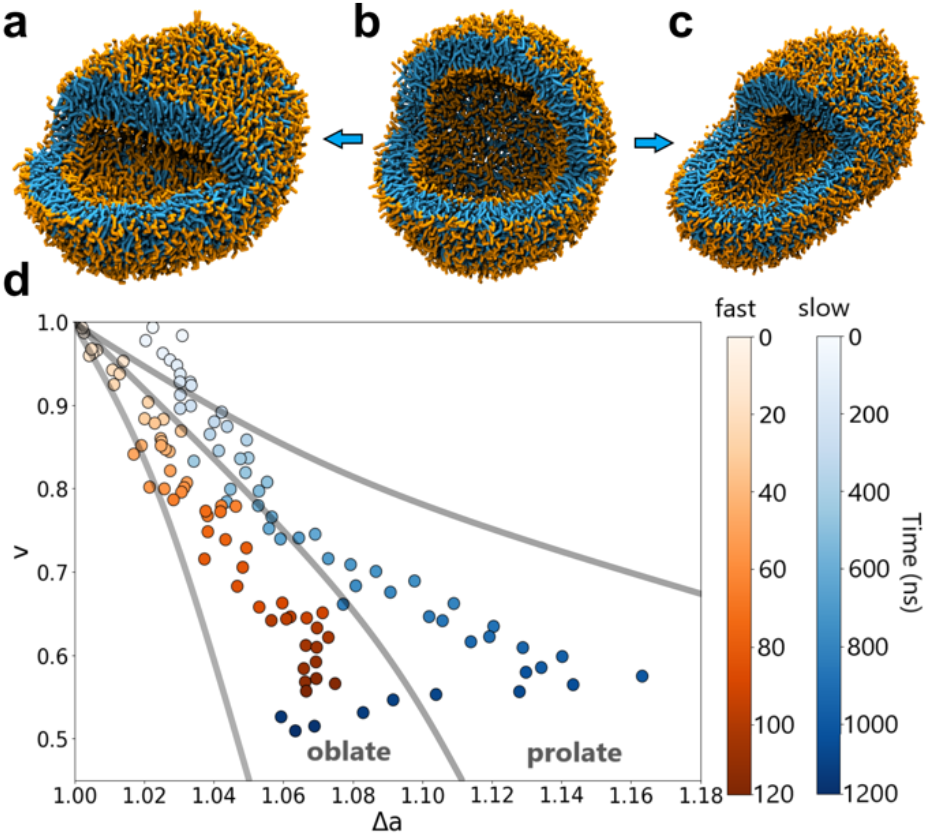
Effect of pumping rate during hypertonic osmotic shock. The snapshots in (a) to (c) show the shape deformation of the initially spherical vesicle (b) after 40% volume reduction using a slow pumping rate of 0.1% (a) and a fast pumping rate of 1% (c). Water molecules are omitted for clarity. (d) Each dot in the shape diagram (relative volume vs. reduced area difference) represents the vesicle shape after a single pumping cycle. The color indicates the temporal evolution for the two pumping rates (white to dark red/blue). The oblate and prolate regions are indicated according to an experimental study by Käs *et al*. [36].

Starting from a spherical vesicle (Figure 2b), we observe vesicle deformations towards a prolate (Figure 2a) and oblate (Figure 2c) shaped vesicle when pumping with a rate of 0.1% and 1%, respectively. To monitor shape changes over time we calculated the reduced area difference Δa and the reduced volume v of the vesicle after each pumping cycle. Since every possible vesicle shape has its unique v/Δa ratio, the shape change over time can easily be visualized in a scakerplot (Figure 2d), where each dot represents the vesicle shape after one pumping cycle along the entire length of the trajectory. The isochores in Figure 2d defining oblate and prolate regions originate from an experimental study by Käs *et al*. [3ti]. For the faster pumping rate (red), the spherical vesicle turns into an oblate one within a couple of pumping cycles (Figure 2c). On the contrary, slow water relocation (blue) preferably leads to prolate vesicles (Figure 2a). However, removing more than 40% of the initial spherical volume causes a sudden change toward an oblate shape, comparable to the shape obtained with the faster pumping rate. In a 5 μs steady-state simulation (Supplementary Figure 1), we observe that the shape of the vesicle that was subjected to the faster pumping rate shows a significant shift over time. The vesicle obtained from the slow-pumping simulation, on the other hand, exhibited a rather constant area difference, indicating that the fast pumping resulted in an accumulation of membrane stress that was relaxed during the steady-state simulation.

### The exploding vesicle

Next, we simulate a hypotonic shock, i.e., water is translocated from the exterior to the interior of a vesicle causing it to expand and eventually to rupture. Here we validate whether Shocker is still able to relocate water particles to the inner compartment when the internal pressure rises. To this end, we created a large simulation box of 60 nm edge length containing a POPC vesicle with an initial radius of approximately 15 nm. To make the system more realistic and add another level of difficulty for the solvent insert process we also added 0.15 mol NaCl. For this simulation a pumping rate of 200 particles every 2 ns was used, corresponding to 0.2% of the initial vesicle volume. After each pumping cycle, a short equilibration run of 20 ps with a timestep of 2 fs is executed, since the rise of internal pressure causes overlap of particles during the insertion step.

To quantify the hypotonic shock, vesicle radius, and membrane thickness were calculated after each pumping cycle. Hereby the thickness is measured as the distance between the glycerol linkers of the lipids. **Figure 3a** shows that the continuous insertion of water molecules leads to swelling of the vesicle. The accompanied rise in internal pressure causes the membrane to expand before it finally ruptures, which is in line with the results of an earlier study [37]. During the pumping process, the vesicle volume grows, causing a linear increase in the radius of the outer and inner leaflet by 2 nm and 2.5 nm, respectively (**Figure 3b**). This difference can be explained by the decreasing membrane thickness as shown in **Figure 3c**. At the rupture point, the vesicle volume increased to approximately 160% of its original volume. If a pore is present at the beginning of a pumping cycle, Shocker detects the non-contiguous membrane structure and switches to a regular (no-pumping) simulation with a default duration of 500 ns.

**Figure 3.**
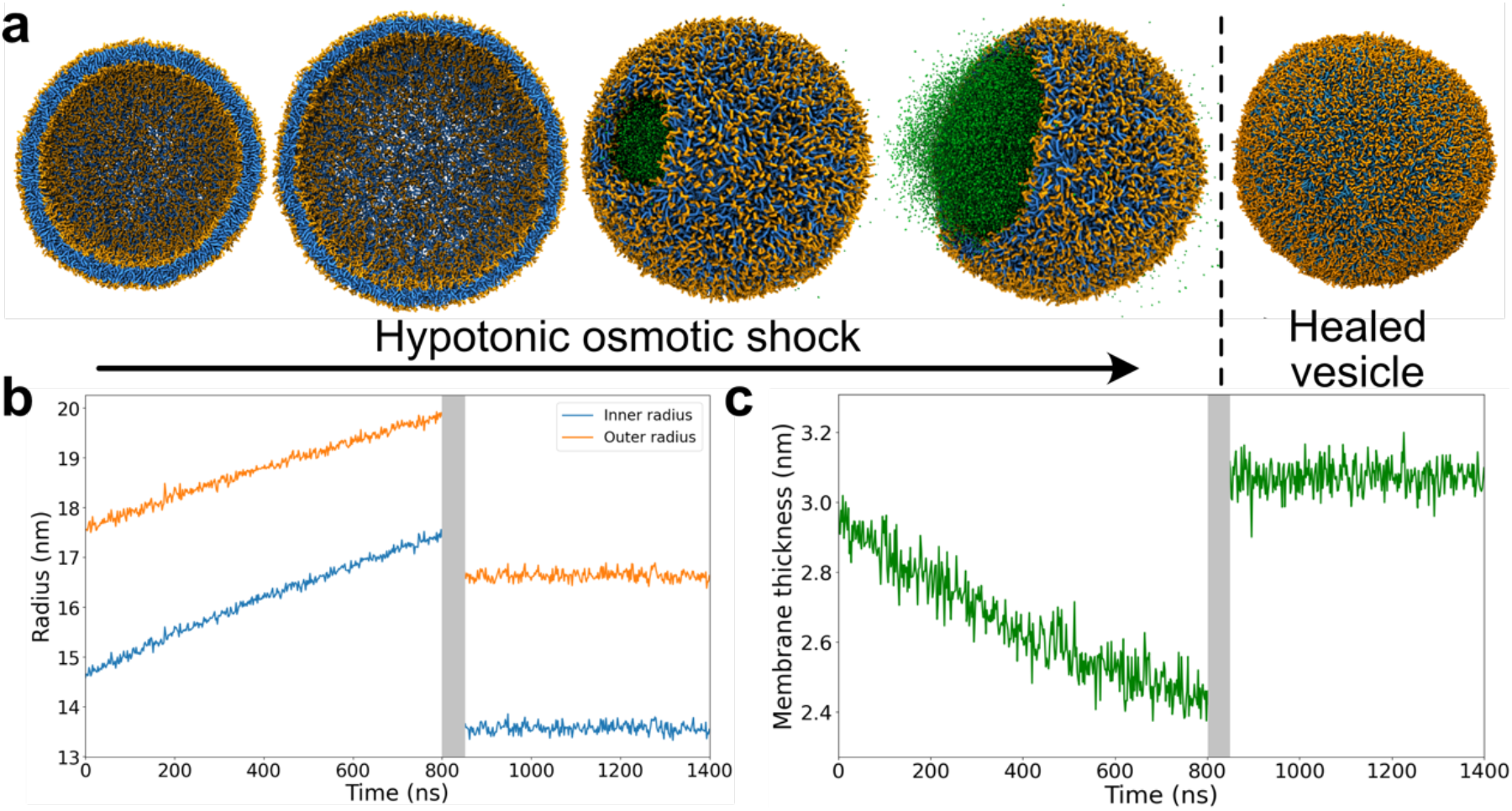
Hypotonic osmotic shock. (a) Snapshots of the stages of a hypotonic shock. The initial stage is an approximately tensionless vesicle filled with water and ions (omitted for clarity). Subsequently, Shocker pumps water from the exterior to the interior compartment, thereby increasing the internal pressure, which leads to thinning of the stressed membrane. Eventually, a pore is formed in the membrane due to the pressure inside. This pore expands fast, releasing excess water (green) and membrane stress. Finally, the vesicle undergoes a rapid healing process and returns to a closed state. The change in (b) radius of the outer and inner leaflet and (c) membrane thickness is shown as a function of simulation ‘me. The gray area in (b) and (c) indicates the presence of a pore.

During a rapid healing period of less than 50 ns, as indicated by a gray bar in **Figure 3b-c**, the excess water particles are released and the membrane returns to a closed state. At the end of the simulation, both radii and the volume of the vesicle are significantly smaller than those of the initial vesicle. Also, the final membrane thickness is slightly larger than the initial thickness. Comparing the number of lipids per leaflet at the beginning of the simulation to the state after the pore closed, we find that the number asymmetry decreased by 8%. This means, that 123 POPC lipids, which were originally located in the outer leaflet, are now part of the inner leaflet. Although the pore was only present for a few nanoseconds, the large rim (**Figure 3a**) allowed a fast exchange of lipids between the two leaflets.

### Deformation of a phase-separated, polymer-filled vesicle

In the third test case, we apply a hypertonic osmotic shock to a phase-separated vesicle with a crowded, polymer-filled inner compartment. We constructed a spherical CG vesicle with an initial radius of 15 nm and a membrane composed of three different lipid types, DPPC/CHOL/DFPC in a 0.3/0.2/0.5 ratio. The membrane was set up in an already phase-separated state, as can be seen in **Figure 4** (top), with the liquid-ordered (L_o_) domain, containing the fully saturated DPPC lipids and CHOL, on the left and the liquid disordered (L_d_) domain, containing the polyunsaturated DFPC lipids, on the right. Subsequently, the vesicle interior was filled with dextran and polyethylene glycol (PEG), which are known to exhibit liquid-liquid phase separation. The dextran and PEG phase are aligned with the L_o_ and L_d_ domain, respectively. This setup roughly resembles the experimental setup presented by Andes-Koback *et al*. [19], who studied asymmetric cell division by applying an osmotic shock on phase-separated and crowded lipid vesicles. As in the previous example, we used a pumping rate of 200 water particles every 2 ns, relocating approximately 0.2% of the initial vesicle solvent volume each pumping cycle. After each pumping cycle, a short equilibration run of 20 ps with a timestep of 2 fs was executed.

**Figure 4.**
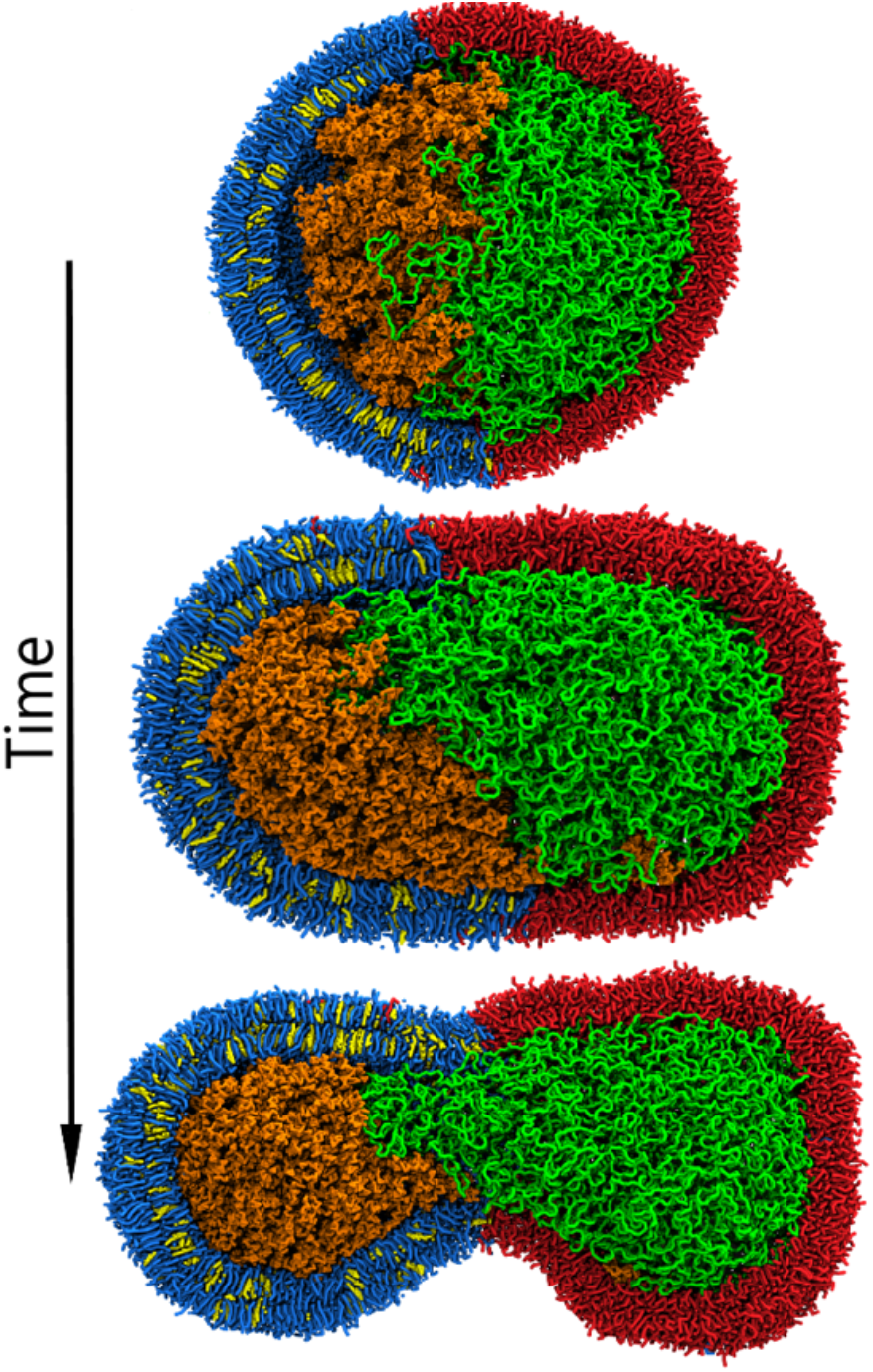
Hypertonic osmotic shock applied to a polymer-filled vesicle. The initial vesicle (top) consists of a phase-separated DPPC/CHOL/DFPC membrane depicted in blue, yellow, and red, respectively, and filled with dextran and PEG (orange and green). During a hypertonic osmotic shock, the vesicle first elongates and forms a prolate (middle) and eventually ends up with a dumbbell shape (bottom) with the neck being located at the interface of the two lipid domains.

The deformation of the initially spherical vesicle over the course of 185 pumping cycles is shown in **Figure 4**. As the volume decreases the vesicle starts to elongate and eventually deform into a dumbbell shape due to the line tension between the two lipid phases. Simultaneously, the phase-separated polymer mixture in the vesicle interior minimizes the interfacial area, further facilitating the formation of a dumbbell-shaped vesicle. This finding is in line with the observation of Andes-Koback *et al*. [19] that these vesicles can split into daughter cells that contain either the PEG or the dextran solution. After a volume reduction of 35%, we continued with a 5 μs steady-state simulation. A detailed shape analysis (see Supplementary Figure 2) revealed that despite the slow pumping rate of 0.2%, the vesicle shape shifted toward an even more prolate state while perfectly maintaining the volume.

### Membrane proteins

So far, we only considered pure lipid membranes in our test cases. With the next system, we want to show that Shocker can also be applied to vesicles with a more complex membrane composition including proteins. To this end, we constructed a CG POPC vesicle with a radius of 1ti nm including 25 aquaporin proteins (**Figure 5a**). Again, the simulation was performed with a pumping rate of 200 particles every 2 ns, corresponding to 0.2% of the initial vesicle volume. For reference, we performed the same simulation for a pure POPC vesicle.

**Figure 5.**
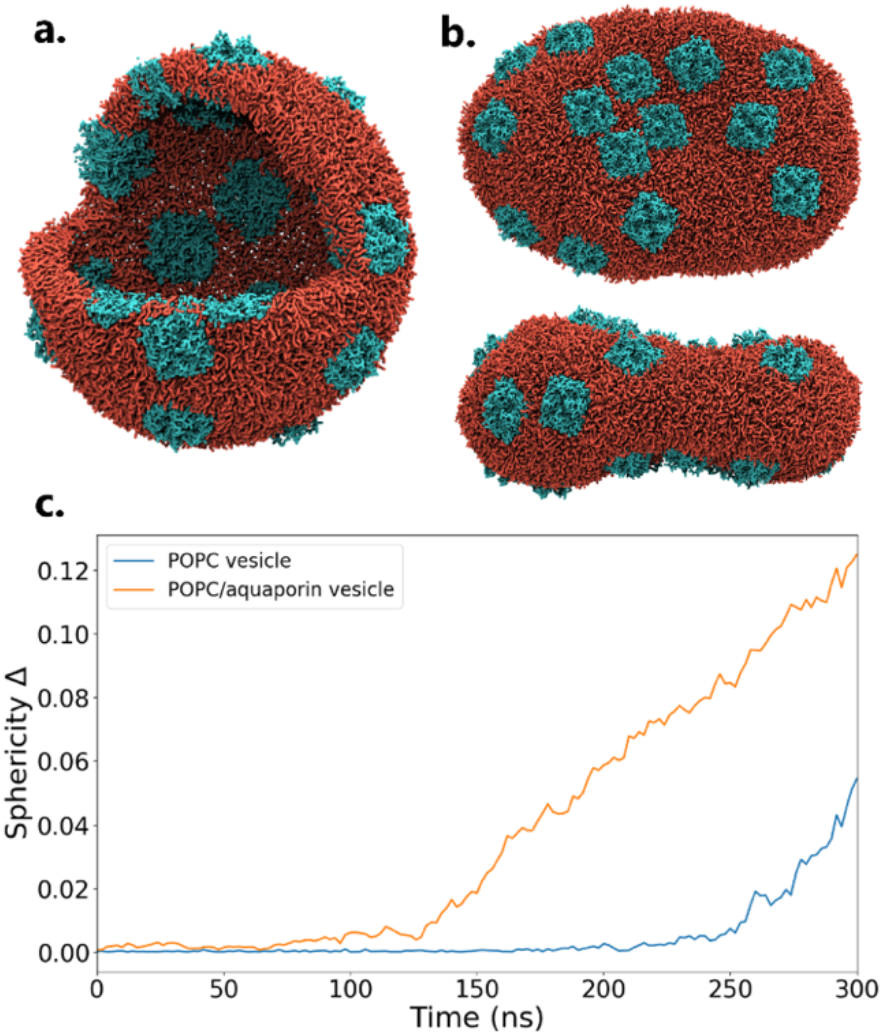
Hypertonic osmotic shock applied to a POPC vesicle with and without transmembrane proteins. Snapshots show (a) the initial spherical vesicle with 25 aquaporins (cyan) embedded in a POPC membrane (red) and (b) the top and side view of the vesicle after a 60% volume reduction. (c) The shape deformation is quantified using the sphericity analysis (described in Supplementary Methods).

**Figure 6.**
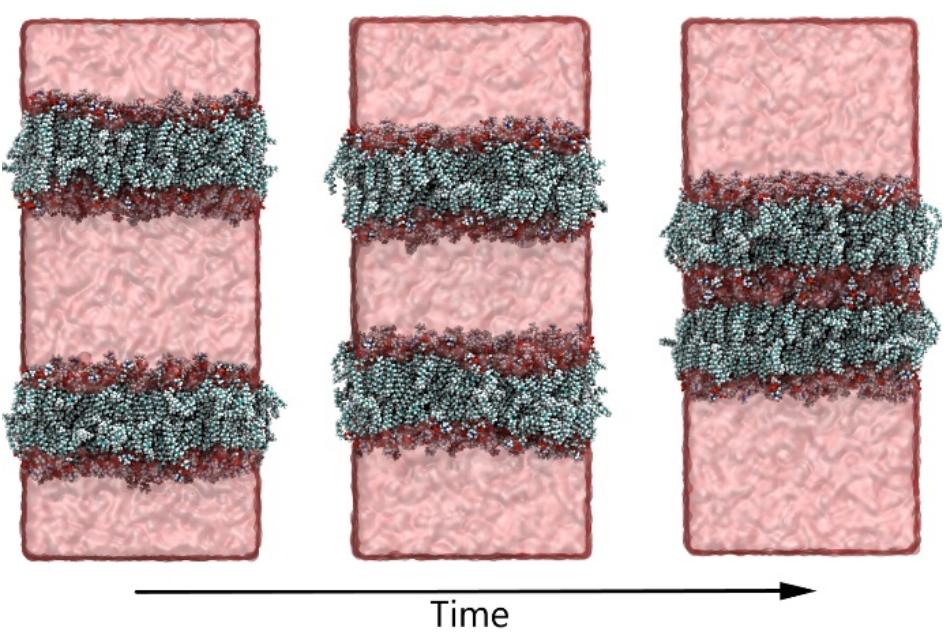
Hypertonic osmotic shock applied to an atomistic double bilayer system. Water is relocated from the central compartment between the bilayers to the space outside the bilayers (crossing the periodic boundaries). Over ‘me the membranes gradually move closer together.

After a volume reduction of 60%, the initially spherical vesicle transformed into an oblate shape with the aquaporins mostly situated on the flat surfaces, as can be seen in the top and side view in **Figure *5*b**. The areas with a strong curvature contained little to no proteins. A video of the deformation process can be found in the Supporting Information.

To quantify the shape changes of the vesicles over the course of the simulation, we computed the sphericity parameter Δ (see Supplementary Methods). A sphericity value of 0 corresponds to a perfect sphere, whereas a value above zero indicates a deviation from a spherical shape. The sphericity analysis in **Figure *5*c** shows the initially spherical vesicle (sphericity parameter close to zero) experiences sudden shape change at around 125 ns in the presence of aquaporin, followed by a steady increase in asphericity. In the case of a POPC vesicle without proteins, however, the shape transformation develops more gradually. During a 5 μs steady-state simulation, the final vesicle of the pumping simulation turned slightly more oblate following the movement of aquaporins toward the flat surfaces of the vesicle. To quantify this, we calculated the oblicity parameter S (see Supplementary Figure 3). The oblicity parameter S becomes increasingly negative, confirming the shift toward an even more oblate-shaped vesicle.

### Atomistic osmotic shock

Lastly, we demonstrate that Shocker is not only compatible with CG but also with AA systems. To reduce the computational costs, we chose a double bilayer system instead of a vesicle. The system consists of two POPC bilayers in a 10×10×25 nm_3_ box, initially positioned with an 8 nm gap between the two membranes. A hypertonic osmotic shock with a pumping rate of 50 water molecules every 2 ns was performed, corresponding to relocating 0.1ti% of the volume of the central compartment each cycle.

Note that Shocker treats the simulation box as non-periodic. This means that in the case of a double bilayer system, three water clusters are found. Since the inner compartment is recognized as such, the remaining clusters are concatenated to end up with one final cluster for the outer compartment. This allows for relocating water molecules evenly to both the upper part and the lower part of the box.

### Performance

To benchmark the performance of Shocker we measured the time needed to complete one pumping cycle. This was done for various test cases with different system sizes ranging from 152.000 to 18.000.000 CG particles, where we relocated 200 solvent particles. Subsequently, we performed tests in which the total number of particles was kept fixed at 750.000, while the number of relocated particles ranged from 100 to 6000. In both cases, a bin-edge size of 1.3 nm was used. All test systems were run on the same Ubuntu desktop computer containing an Intel Xeon CPU (ti cores, hyper-threading, 4.5 GHz) and 1tiGB of DDR4 memory. The results of these test runs are collected in Figure 7.

**Figure 7.**
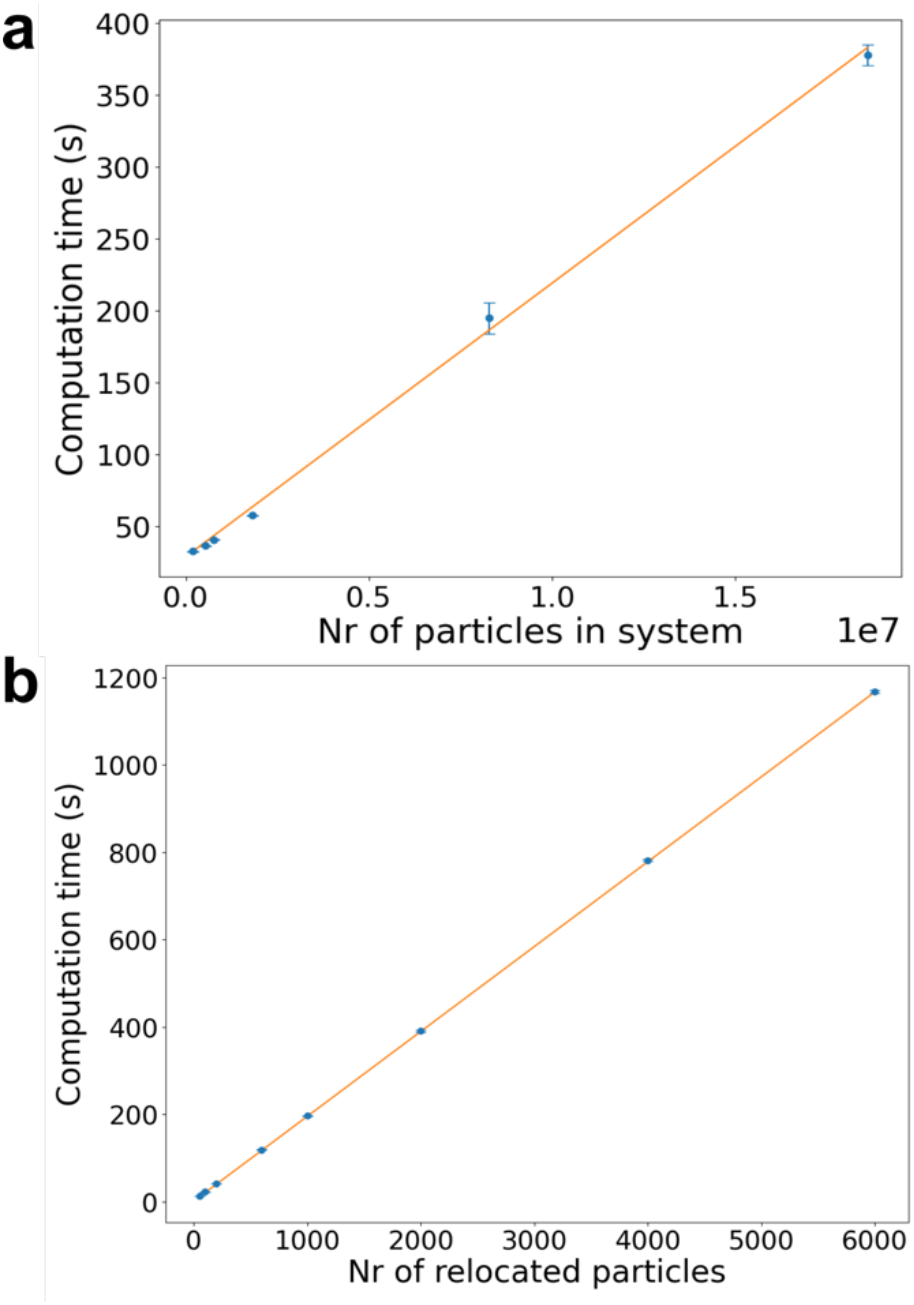
Computation time of a single pumping cycle. The computation time of a single pumping cycle is measured for(a)different system sizes when relocating 200 particles and for (b)different numbers of particles to be relocated in a fixed system size of 750.000 particles. Error bars indicate the standard deviation obtained from 10 replicas.

**Figure 7a** shows that there is a linear relationship between the time Shocker needs to perform one pumping cycle and the number of particles the systemconsists of. When relocating 200 particles Shocker is able to complete one pumping cycle within 6 minutes for all tested system sizes. **Figure 7b** shows that there also exists a linear relationship between the number of relocations and computation time for one cycle. For a system size of 750.000 CG particles, which is around the size of most of the systems in this study, Shocker needs 42 seconds to perform one relocation cycle. However, when relocating 6000 particles, which is almost 1% of the content of the box, Shocker needs 20 minutes of computation time. Considering that for a system of this size, a single 2 ns production run takes 18 minutes even this relocation time is acceptable.

## Discussion & Conclusion

Osmosis is a fundamental process that is coupled to many processes at the cellular level [38]. Beyond being a simple environmental factor, osmotic shocks are also frequently used as a tool in synthetic chemical biology to, for example, manipulate membrane shapes of cells [11, 39]. Despite the widespread occurrence in biology and synthetic biology, a molecular-level picture of how membrane shape deformations are steered by osmosis remains hidden from experimental techniques and still poses many open questions. To answer those questions and elucidate the molecular driving forces, simulation approaches such as coarse-grained molecular dynamics simulations are commonly employed [23, 40, 41] Since simulating an osmotic shock explicitly in MD is a time-consuming process, researchers circumvent it by either using pre-deformed vesicles [26] or removing solvent molecules from the interior in the initial setup [24]. Even though these studies can give valuable insights into vesicle shape transformations including budding, fusion, and fission, they allow very little control over the final membrane shapes or the deformation pathways. To allow the efficient and controlled simulation of osmotic shocks in MD using explicit water models, we presented and benchmarked the Shocker Python program.

### Robust Simulation Protocol

Shocker mimics an osmotic shock by moving solvent particles between compartments, according to the direction of the osmotic shock, in several pumping cycles. A pumping cycle starts by identifying an inner and outer solvent compartment using an efficient binning algorithm. Subsequently, a number of solvent molecules defined by the pumping rate is moved to the other compartment depending on the type of osmotic shock (hypertonic or hypotonic). Instead of just randomly placing solvent molecules, it is taken care that the system is perturbed as little as possible by finding the largest gap within a solvent bin. Finally, a short equilibration is performed before a new cycle begins. We note that the program not only implements hypertonic and hypotonic shock but additionally is capable of performing a steady-state simulation that keeps the number of solvent molecules constant by counteracting the natural solvent permeation.

To assess the extent of perturbation due to the solvent relocation we conducted two benchmark osmotic shocks utilizing a slow and fast pumping rate. The test system consisted of a water box at the all-atom and the CG Martini level, in which either 0.05% or 1% of water molecules were relocated within the water box. The distance distribution between the newly placed water particles and all other water particles in Supplementary Figure 4 showed that the method produces no direct overlaps even for the fast pumping rate. This conclusion was further supported by the fact the simulation can be continued after relocating the water particles without the need for energy minimization. While the relocation does produce a sudden increase in potential energy, this energy is quickly dissipated (see Supplementary Figure 5). We note that in general within the Shocker protocol velocities are preserved after water relocation even for the more complex test cases. However, especially in crowded systems, water molecules may have to be removed from the immediate solvation shell of other molecules such as proteins and polymers in solution. To ensure a stable simulation protocol, Shocker offers the possibility to automatically add energy minimization and equilibration steps into the pipeline even though for most systems they are not required. Only the crowded polymer-filled vesicles required such additional measures from the test cases surveyed. Additionally, as a built-in safety net against numerical failure of simulations, Shocker returns to a state before the errors occurred and tries to relocate other water particles to new positions. This continues until the production run finishes correctly.

### Realistic Membrane Shapes

To assess that Shocker produces realistic membrane shapes we tested the protocol on increasingly complex membrane systems. First, we applied a hypertonic osmotic shock to a simple POPC vesicle and examined the effect of different solvent pumping rates on its shape. To monitor the shapes over time we constructed a shape phase diagram, which clearly showed two distinct shape pathways generally obeying the experimentally determined shape boundaries [3ti]. Slower pumping rates result in more prolate shapes as opposed to faster pumping rates, which yield oblate shapes. This finding is consistent with an earlier simulation study [30]. After the target volume was reached, a 5 μs steady-state simulation was performed on both vesicles. While keeping the interior vesicle volume constant, membrane shape changes could be observed. Slower pumping rates yield more stable shapes and are closer to equilibrium shape deformations, while the vesicle subjected to a faster pumping rate experienced a significant shift in shape over time. This example demonstrates that the user needs to carefully choose an appropriate pumping rate. We recommend 0.2% as a good compromise of being fast while allowing sufficient relaxation time to avoid kinetic artifacts.

By applying a hypotonic osmotic shock to a POPC vesicle, we demonstrated that our protocol can be used to inflate a vesicle up to the point of rupture. Furthermore, after rupture the program automatically switches to a regular no-pumping simulation, allowing the vesicle to heal. An interesting result was the smaller radius and volume as well as the larger membrane thickness of the healed vesicle compared to the initial vesicle. For this observation, we consider two possible explanations. On the one hand, it is possible that the initial vesicle is not completely tensionless. In this case, the vesicle initially contains a larger amount of water than energetically favorable and/or has an imbalance of lipids between the inner and outer vesicle leaflet. This can be corrected by the efflux of water after rupture and by accelerated lipid flip-flop at the large pore rim, which results in a vesicle with a slightly smaller radius than the initial one. On the other hand, with the rapid opening followed by a fast healing process, more water might be excreted than inserted during the osmotic shock. This kinetic effect would also result in a vesicle with a smaller volume than the initial one. While further analysis is needed to confirm either hypothesis, we have no reason to believe the observed result is an artifact of the osmotic shocking protocol.

To further increase complexity, we considered a system consisting of a phase-separated membrane with an aqueous two-phase polymer system on the interior. Previously, similar systems have been shown to undergo asymmetric fission in experiments under osmotic shock [19]. While we did not see fission, the system deforms from a spherical vesicle to a pre-fission, dumbbell state. The polymer interior remains phase separated and aligns with the neck at the interface of the two lipid phases. This behavior is consistent with the experimental results observed for similar systems [19, 42, 43].

Next, we tested membrane deformations in the presence of proteins. It is well known that proteins induce membrane curvature and have an intrinsic curvature preference [44]. Given the crowdedness of realistic cell membranes, it is of paramount importance to capture shape deformations in the presence of proteins. Our test system consisted of a vesicle with 25 aquaporins embedded into a POPC membrane. Under a hypertonic shock with a volume reduction of 40% the membrane adopts a flat disk shape. The strongest curvature occurs in the areas with the lowest protein density. The difference in shape development between a pure POPC vesicle and the protein-containing vesicle must therefore originate from the proteins embedded in the membrane, which rigidify the membrane. A non-spherical shape can alleviate the energetic cost of bending such a stiff membrane, in particular as the system can form protein-rich and protein-poor domains. The vesicle, therefore, remains spherical until a point where the low internal pressure makes the vesicle collapse into a flat disk shape. Hereby, the strong curvature appears in the most flexible membrane areas, i.e., those with the lowest protein density. The early deformation in the protein-containing vesicle can be explained by the fact that these membrane proteins prefer a flat membrane surface and thus promote an early deformation to create flat surfaces. This phenomenon became even more clear when subjecting the final vesicle to a 5 μs steady-state simulation, where the proteins moved to flat membrane areas, resulting in a more oblate-shaped vesicle. Given that aquaporin has previously been reported to prefer zero curvature regions on the membrane [45], these results validate the applicability of our protocol to membranes containing proteins.

Lastly, we tested Shocker on an AA system consisting of two flat POPC bilayers. A hypertonic shock was performed, therefore reducing the volume of the compartment between the bilayers. As expected, the bilayers moved closer together consistent with a hypertonic shock.

### Limitations & Future Extensions

To measure the performance of the water relocation code we benchmarked systems ranging from 150.000 to 18 million particles. On a desktop computer, a single pumping cycle is completed within minutes when relocating 200 particles. Generally, the relocation is therefore much faster than any equilibration and production run on large systems. Hence, we consider Shocker performant enough even for the largest currently studied systems. In principle, the speed of water relocation can even be increased by increasing the number of relocated water particles. However, as mentioned earlier this can influence the shape deformation pathways and potentially cause numerical stability problems.

An important current limitation of the tool is that Shocker only works with two water compartments. Even though this is sufficient for applications to vesicles, tubes, and double bilayer systems, many higher-order membrane shapes of interest, such as multi-lamellar vesicles, exist. The principal limitation to two compartments results from the binning procedure used to identify the compartments. Combining Shocker with more advanced protocols such as the MDVoxel Segmentation protocol [4ti] could be a future extension to make such systems possible.

Finally, we note that Shocker implements some common shape analysis metrics which can be used to characterize the vesicle shapes along the pumping simulations. In order to compute properties such as the area and volume of the deformed vesicles, first any periodic boundary crossings need to be removed to generate a continuous surface required in the second step. The surface of each leaflet is then generated using the pyvista surface reconstruction method [47] which allows an estimation of the vesicle area and volume. Removing periodic boundary crossings using the in-house method, however, can fail if the object crosses the PBC multiple times. If the reconstruction fails, no continuous surface can be generated, which ultimately leads to a failure of the analysis. Using more advanced algorithms such as MDVWhole [48] would be an option to make this step more robust and ensure the analysis protocol works for shapes split across the PBC in arbitrary ways. The shape reconstruction can also fail when the membrane displays regions of high curvature where the separation between leaflets is so little that the surfaces cannot be reconstructed properly by the used algorithm. Further surface refinement might be required to proceed with the analysis.

We conclude that the Shocker Python program outlined in this paper offers a robust protocol to study the effect of osmotic shocks on membranes in a realistic and direct manner.

## Methods

### System setup

All CG vesicle systems used in this work were constructed using TS2CG [49], an MD tool designed to generate initial configurations for lipid membrane structures with user-defined compositions (including proteins) and shapes. Lipid parameters were taken from the recently published Martini 3 [34] force field. Topology and structure files of CG proteins were generated with Martinize2 [50] also using the Martini 3 [34] forcefield. Elastic bonds were applied to aquaporin (PBD ID: 1J4N [51]) to maintain the secondary structure as well as the tetrameric assembly. To fill the interior of those vesicles with polymers Polyply [52] was used, a Python suite designed for generating input files and system coordinates for macromolecules. Topologies for PEG and dextran were taken from Polyply [52] and our recently published Martini 3 extension for carbohydrates [53], respectively. System solvation and placement of sodium chloride ions (if needed) were done using the solvation script provided by TS2CG [49].

The AA double bilayer system was constructed in two steps: First, we generated a single POPC bilayer using the CHARMM-GUI [54] with CHARMM3ti forcefield [35] parameters and TIP3P water. Subsequently, Gromacs tools were used to duplicate the entire system in the z-dimension.

### System details

Supplementary Table 1 displays the system size and composition of all the setups used in this work. The lipid names are consistent with the names used in the Martini CG lipidomics database (hkp://cgmartini.nl/index.php/force-field-parameters/lipids). The lipid ratio in Supplementary Table 1 is determined by TS2CG [49] during lipid placement.

After placing the lipids and initial energy minimization using the steepest descent algorithm, a series of equilibration runs with different membrane pore sizes was performed (as described in a previous work [55]). This allows for both lipid redistribution between the two leaflets and equilibration of the solvent content between the interior and exterior solvent compartment. During the equilibration, the pores were gradually closed before we continued with the production run.

### Simulation parameters

All simulations were executed with GROMACS version 2021.2 [5ti]. Simulation parameters for CG simulations were based on standard values used in the Martini benchmarking paper [57]. CG simulations were performed in the NPT ensemble at a constant pressure and temperature of 1 bar and 310 K, respectively, except for the simulation with the polymer-filled vesicle, which was performed at a temperature of 323 K. The pressure and temperature were maintained using the anisotropic Berendsen coupling [58] and velocity rescaling thermostat [59], respectively. Furthermore, for production runs an integration time step of 20 fs was chosen using a leap-frog algorithm for integrating the equations of motion.

For the AA simulation, the pressure of 1 bar and temperature of 310 K were maintained using the Parrinello-Rahman pressure coupling [60] and the Nose-Hoover thermostat [ti1], respectively. An integration step of 2 fs was used.

### Analysis details

All videos and snapshots of the simulations were rendered with VMD [ti2]. Calculation of the volume, area, and sphericity parameter of vesicles is an intrinsic feature of Shocker, as explained in more detail in the Supplementary Information.

### Use and availability

Shocker can be found in our GitHub repository (hkps://github.com/marrink-lab/shocker) and installed according to the README file. It is accompanied by several tutorials and datasets to perform hypertonic and hypotonic osmotic shock simulations. The initial configurations of all experiments described in this paper, as well as the corresponding command line options to invoke Shocker, can also be found on the GitHub page.

## Acknowledgments

We would like to thank the Center for Information Technology of the University of Groningen for their support and for providing access to the Peregrine/Hábrók high-performance computing cluster and SURFsara for providing access to Snellius. S.J.M. acknowledges funding from the ERC *via* an Advanced grant “COMP-O-CELL”.

## Notes

### Competing Interest Statement

The authors have declared no competing interest.

## References

1. Scharwies, J.D. and J.R. Dinneny, Water transport, perception, and response in plants. Journal of Plant Research, 2019. 132: p. 311–324.

2. Csonka, L.N., Physiological and genetic responses of bacteria to osmotic stress. Microbiological Reviews, 1989. 53: p. 121–147.

3. Haswell, Elizabeth S., R. Phillips, and Douglas C. Rees, Mechanosensitive Channels: What Can They Do and How Do They Do It? Structure, 2011. 19: p. 1356–1369.

4. Berika, M., M.E. Elgayyar, and A.H.K. El-Hashash, Asymmetric cell division of stem cells in the lung and other systems. Frontiers in Cell and Developmental Biology, 2014. 2.

5. Sunchu, B. and C. Cabernard, Principles and mechanisms of asymmetric cell division. Development, 2020. 147.

6. Exterkate, M. and A.J.M. Driessen, Synthetic Minimal Cell: Self-Reproduction of the Boundary Layer. ACS Omega, 2019. 4: p. 5293–5303.

7. Olivi, L., et al., Towards a synthetic cell cycle. Nature communications, 2021. 12(1): p. 4531.

8. Miele, Y., et al., Shape Deformation, Budding and Division of Giant Vesicles and Artificial Cells: A Review. Life, 2022. 12: p. 841.

9. Snead, W.T., et al., Membrane fission by protein crowding. Proceedings of the National Academy of Sciences, 2017. 114.

10. Allain, J.M. and M.B. Amar, Budding and fission of a multiphase vesicle. The European Physical Journal E, 2006. 20: p. 409–420.

11. Steinkühler, J., et al., Controlled division of cell-sized vesicles by low densities of membrane-bound proteins. Nature Communications, 2020. 11: p. 905.

12. Inaoka, Y. and M. Yamazaki, Vesicle Fission of Giant Unilamellar Vesicles of Liquid-Ordered-Phase Membranes Induced by Amphiphiles with a Single Long Hydrocarbon Chain. Langmuir, 2007. 23: p. 720–728.

13. Baldauf, L., et al., Actomyosin-Driven Division of a Synthetic Cell. ACS Synthetic Biology, 2022. 11: p. 3120–3133.

14. Kohyama, S., A. Merino-Salomón, and P. Schwille, In vitro assembly, positioning and contraction of a division ring in minimal cells. Nature Communications, 2022. 13: p. 6098.

15. Stachowiak, J.C., et al., Membrane bending by protein–protein crowding. Nature cell biology, 2012. 14(9): p. 944–949.

16. Miele, Y., et al., Self-division of giant vesicles driven by an internal enzymatic reaction. Chemical Science, 2020. 11: p. 3228–3235.

17. Farge, E. and P.F. Devaux, Shape changes of giant liposomes induced by an asymmetric transmembrane distribution of phospholipids. Biophysical Journal, 1992. 61: p. 347–357.

18. De Franceschi, N., et al., Synthetic membrane shaper for controlled liposome deformation. ACS nano, 2022. 17(2): p. 966–978.

19. Andes-Koback, M. and C.D. Keating, Complete Budding and Asymmetric Division of Primitive Model Cells To Produce Daughter Vesicles with Different Interior and Membrane Compositions. Journal of the American Chemical Society, 2011. 133: p. 9545–9555.

20. Dreher, Y., et al., Division and regrowth of phase-separated giant unilamellar vesicles. Angewandte Chemie International Edition, 2021. 60(19): p. 10661–10669.

21. Sakuma, Y. and M. Imai, Model System of Self-Reproducing Vesicles. Physical Review Lekers, 2011. 107: p. 198101.

22. Deshpande, S., et al., Mechanical Division of Cell-Sized Liposomes. ACS Nano, 2018. 12: p. 2560–2568.

23. Pezeshkian, W. and S.J. Marrink, Simulating realistic membrane shapes. Current Opinion in Cell Biology, 2021. 71: p. 103–111.

24. Ghosh, R., et al., Spherical Nanovesicles Transform into a Multitude of Nonspherical Shapes. Nano Lekers, 2019. 19: p. 7703–7711.

25. Markvoort, A.J. and S.J. Marrink, Lipid acrobatics in the membrane fusion arena. Current Topics in Membranes, 2011. 68: p. 259–294.

26. Markvoort, A.J., et al., Lipid-Based Mechanisms for Vesicle Fission. The Journal of Physical Chemistry B, 2007. 111: p. 5719–5725.

27. Markvoort, A.J., R.A. van Santen, and P.A.J. Hilbers, Vesicle Shapes from Molecular Dynamics Simulations. The Journal of Physical Chemistry B, 2006. 110: p. 22780–22785.

28. Kawamoto, S., M.L. Klein, and W. Shinoda, Coarse-grained molecular dynamics study of membrane fusion: Curvature effects on free energy barriers along the stalk mechanism. The Journal of Chemical Physics, 2015. 143(24).

29. Hong, C., D.P. Tieleman, and Y. Wang, Microsecond Molecular Dynamics Simulations of Lipid Mixing. Langmuir, 2014. 30: p. 11993–12001.

30. Yuan, H., C. Huang, and S. Zhang, Dynamic shape transformations of ?uid vesicles. SoG Maker, 2010. 6: p. 4571.

31. Harris, B., G.-y. Liu, and R. Faller, GenEvaPa: A generic evaporation package for modeling evaporation in molecular dynamics simulations. Computer Physics Communications, 2023. 282: p. 108539.

32. Zhuang, Y., et al., Implementation of Telescoping Boxes in Adaptive Steered Molecular Dynamics. Journal of Chemical Theory and Computation, 2022. 18: p. 4649–4659.

33. Guo, J., et al., An Efficient Voxel-Based Segmentation Algorithm Based on Hierarchical Clustering to Extract LIDAR Power Equipment Data in Transformer Substations. IEEE Access, 2020. 8: p. 227482–227496.

34. Souza, P.C.T., et al., Martini 3: a general purpose force field for coarse-grained molecular dynamics. Nature Methods, 2021. 18: p. 382–388.

35. Vanommeslaeghe, K., et al., CHARMM general force field: A force field for drug-like molecules compatible with the CHARMM allatom additive biological force fields. Journal of computational chemistry, 2010. 31(4): p. 671–690.

36. Käs, J. and E. Sackmann, Shape transitions and shape stability of giant phospholipid vesicles in pure water induced by area-to-volume changes. Biophysical Journal, 1991. 60: p. 825–844.

37. Louhivuori, M., et al., Release of content through mechano-sensitive gates in pressurized liposomes. Proceedings of the National Academy of Sciences, 2010. 107: p. 19856–19860.

38. McManus, M.L., K.B. Churchwell, and K. Strange, Regulation of cell volume in health and disease. New England Journal of Medicine, 1995. 333(19): p. 1260–1267.

39. Liu, X., et al., Vesicles balance osmotic stress with bending energy that can be released to form daughter vesicles. The Journal of Physical Chemistry Lekers, 2022. 13(2): p. 498–507.

40. Marrink, S.J., et al., Computational modeling of realistic cell membranes. Chemical reviews, 2019. 119(9): p. 6184–6226.

41. Marrink, S.J., et al., Two decades of Martini: Be[er beads, broader scope. Wiley Interdisciplinary Reviews: Computational Molecular Science, 2023. 13(1): p. e1620.

42. Cans, A.-S., M. Andes-Koback, and C.D. Keating, Positioning lipid membrane domains in giant vesicles by micro-organization of aqueous cytoplasm mimic. Journal of the American Chemical Society, 2008. 130(23): p. 7400–7406.

43. Su, W.-C., et al., Kinetic control of shape deformations and membrane phase separation inside giant vesicles. Nature Chemistry, 2023: p. 1–9.

44. Zimmerberg, J. and M.M. Kozlov, How proteins produce cellular membrane curvature. Nature reviews Molecular cell biology, 2006. 7(1): p. 9–19.

45. Tieleman, D., et al., Insights into lipid-protein interactions from computer simulations. Biophysical Reviews, 2021: p. 1–9.

46. Bruininks, B.M., et al., Sequential voxel-based lea?et segmentation of complex lipid morphologies. Journal of Chemical Theory and Computation, 2021. 17(12): p. 7873–7885.

47. Sullivan, C. and A. Kaszynski, PyVista: 3D plo]ng and mesh analysis through a streamlined interface for the Visualization Toolkit (VTK). Journal of Open Source Software, 2019. 4(37): p. 1450.

48. Bruininks, B.M., T.A. Wassenaar, and I. Vakulainen, Unbreaking Assemblies in Molecular Simulations with Periodic Boundaries. Journal of chemical information and modeling, 2023.

49. Pezeshkian, W., et al., Backmapping triangulated surfaces to coarse-grained membrane models. Nature communications, 2020. 11: p. 1–9.

50. Kroon, P.C., et al., Martinize2 and vermouth: Unified framework for topology generation. arXiv preprint 2212.01191, 2022.

51. Sui, H., et al., Structural basis of water-specific transport through the AQP1 water channel. Nature, 2001. 414(6866): p. 872–878.

52. Grünewald, F., et al., Polyply; a python suite for facilitating simulations of macromolecules and nanomaterials. Nature Communications, 2022. 13: p. 68.

53. Grünewald, F., et al., Martini 3 coarse-grained force field for carbohydrates. Journal of Chemical Theory and Computation, 2022. 18(12): p. 7555–7569.

54. Jo, S., et al., CHARMM-GUI: a web-based graphical user interface for CHARMM. Journal of computational chemistry, 2008. 29(11): p. 1859–1865.

55. Risselada, H.J., A.E. Mark, and S.J. Marrink, Application of mean field boundary potentials in simulations of lipid vesicles. The Journal of Physical Chemistry B, 2008. 112(25): p. 7438–7447.

56. Abraham, M.J., et al., Gromacs: High performance molecular simulations through mul1-level parallelism from laptops to supercomputers. SoftwareX, 2015. 1-2: p. 19–25.

57. De Jong, D.H., et al., Martini straight: Boosting performance using a shorter cutoff and GPUs. Computer Physics Communications, 2016. 199: p. 1–7.

58. Berendsen, H.J.C., et al., Molecular dynamics with coupling to an external bath. The Journal of Chemical Physics, 1984. 81: p. 3684–3690.

59. Bussi, G., D. Donadio, and M. Parrinello, Canonical sampling through velocity rescaling. The Journal of Chemical Physics, 2007. 126: p. 014101.

60. Parrinello, M. and A. Rahman, Polymorphic transitions in single crystals: A new molecular dynamics method. Journal of Applied Physics, 1981. 52: p. 7182–7190.

61. Nosé, S., A unified formulation of the constant temperature molecular dynamics methods. The Journal of Chemical Physics, 1984. 81: p. 511–519.

62. Humphrey, W., A. Dalke, and K. Schulten, VMD: Visual molecular dynamics. Journal of Molecular Graphics, 1996. 14: p. 33–38.

