## Supplementary Information for "Shocker - a molecular dynamics protocol and tool for accelerating and analyzing the effects of osmotic shocks"

#### Supplementary Figures

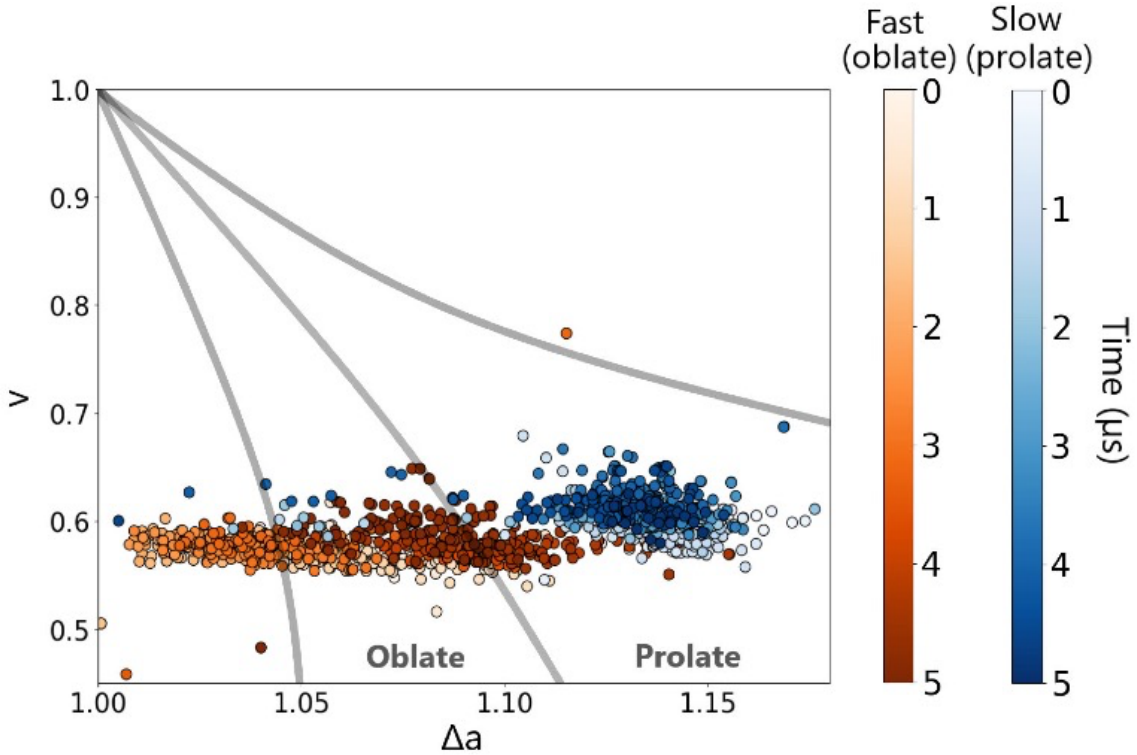


**Supplementary Figure 1 Shape analysis of steady-state simulations of different pumping rates.** A hypertonic shock was performed on POPC vesicles using pumping rates of 0.1% (slow) and 1% (fast) of the initial interior vesicle volume. After a 40% volume reduction, a 5 µs steady-state simulation was launched and the changes in shape for the two pumping rates are indicated. Each dot in the shape diagram represents the vesicle shape after 20 ns, corresponding to 10 evaluations of the vesicle volume. The color indicates the temporal evolution for the two pumping rates (white to dark red/blue).

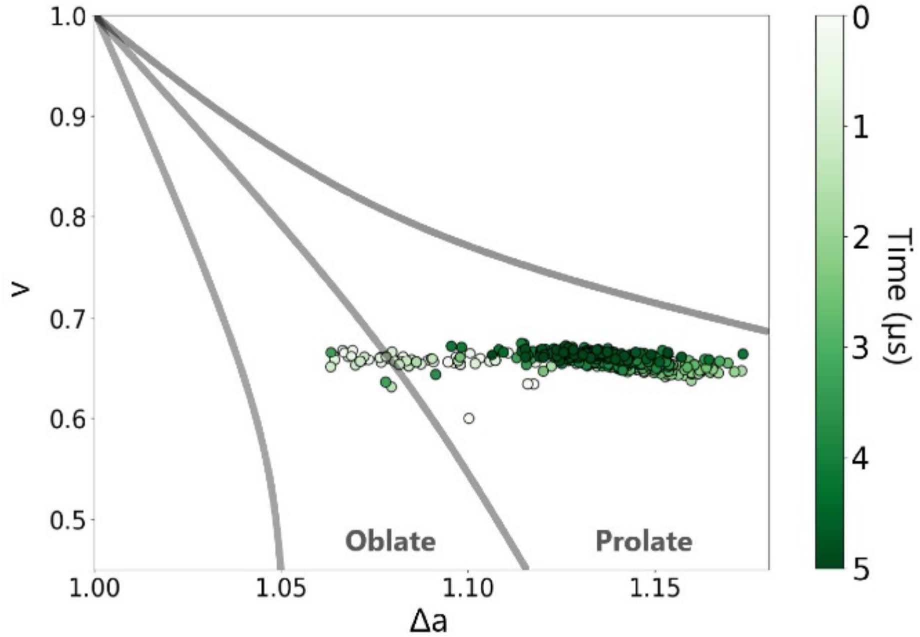


**Supplementary Figure 2 Shape analysis of steady-state simulations for polymer-filled phase-separated vesicle.** A hypertonic shock was performed on a vesicle with a phase-separated DPPC/CHOL/DFPC membrane and filled with dextran and PEG. After a 35% volume reduction of the initial water enclosed water volume, a 5 µs steady-state simulation was launched and the changes in shape are indicated. The color indicates the temporal evolution from white to dark green.

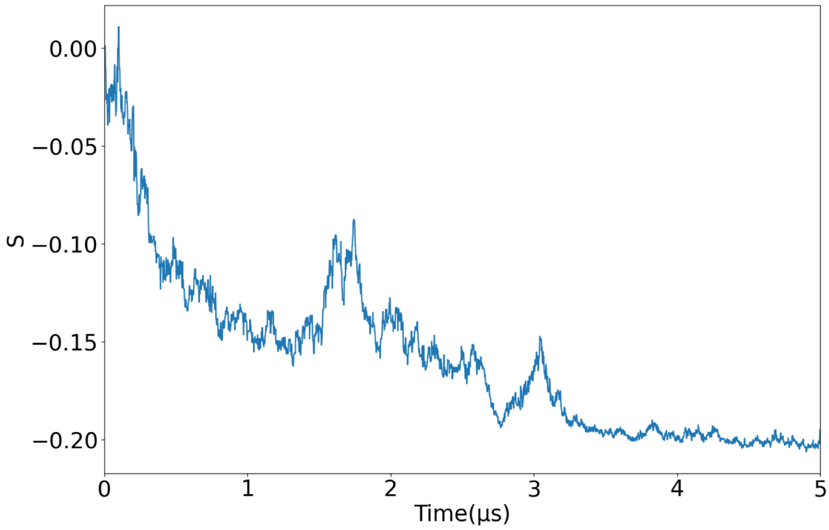


**Supplementary Figure 3 Oblicity analysis of steady-state simulations with and without transmembrane proteins.** A hypertonic shock was performed on a POPC vesicle with and without aquaporin embedded into the membrane. After a 40% volume reduction, a 5 µs steady-state simulation was launched and the changes in oblicity S are calculated as described in the Supplementary Methods. A negative oblicity value indicates an oblate shape.

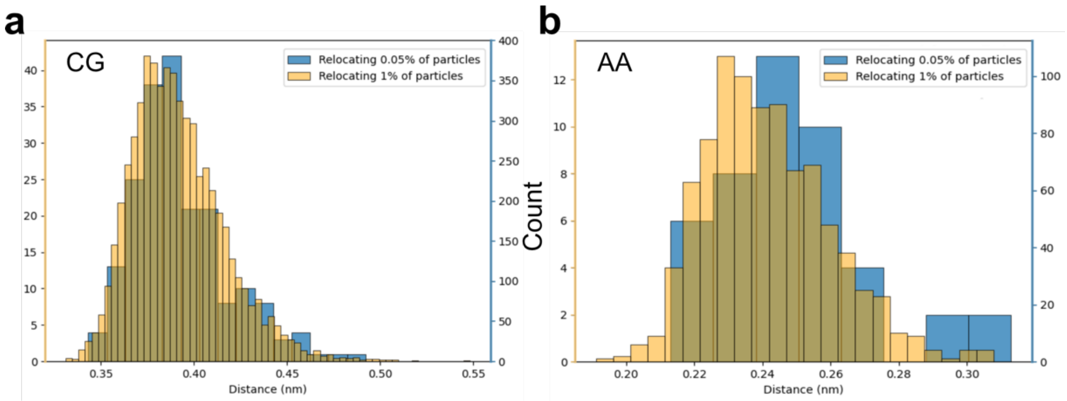


**Supplementary Figure 4 Histograms of water-water distances after relocation.** The center of mass distances of all relocated water particles to the closest other water particles in the bin for (a) a CG system and (b) an AA system. For both systems, we applied pumping rates of 0.05% (yellow) and 1% (blue), corresponding to 300 and 6000 water beads for the CG system or 30 and 600 water molecules for the AA system. The setup is a water box in which particles were moved from one side to the other.

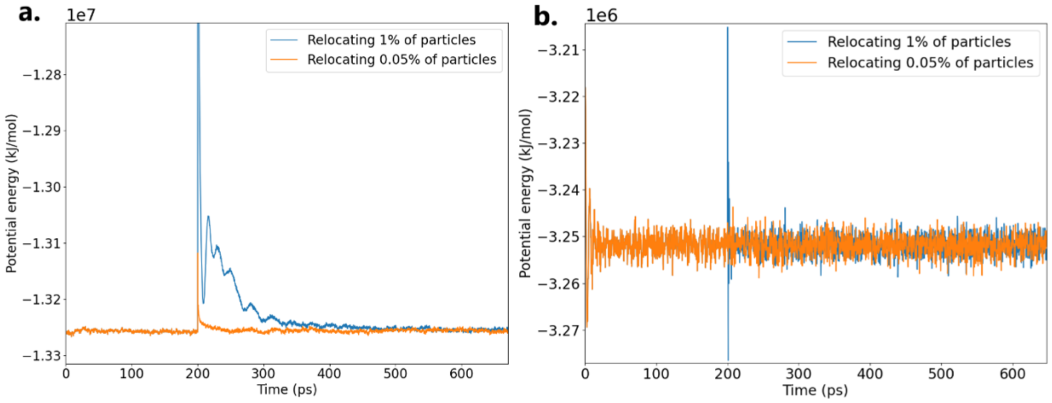


**Supplementary Figure 5 Evolution of potential energy during a pumping cycle.** Water particles were after 200 ps of simulation time for (a) a CG system and (b) an AA system as described in **Supplementary Figure 4**.

#### Supplementary Tables

**Supplementary Table 1 System composition and simulation details for all example systems.** Shocker was used to perform hypertonic and hypotonic osmotic shock simulations for a variety of different systems. Simulation times varied between 200 ns and 2400 ns depending on the pumping rate.

| | Example system | System size (nm) | Membrane composition | Lipid ratio (outer/inner) | Vesicle radius (nm) | Pumping rate (per 2 ns) | Initial interior volume (no. of solvent particles) |
|---|---|---|---|---|---|---|---|
| CG MARTINI | Test pumping rate (hypertonic) | 20x20x20 | POPC | 1797/603 | 7.5 | Slow: 10<br>Fast: 100 | 10 000 |
| CG MARTINI | Exploding vesicle (hypotonic) | 60x60x60 | POPC | 4892/3492[a]<br>4769/3615[b] | 15.9 | 200 | 120 000 |
| CG MARTINI | Polymer vesicle (hypertonic) | 40x40x40 | DPPC/CHOL/DFPC (0.30/0.20/0.50) | 5267/3251 | 16.4 | 200 | 94 000 |
| CG MARTINI | Protein vesicle (hypertonic) | 60x60x60 | POPC with 25 aquaporin (PDB-ID: 1J4N) | 3722/2635 | 15.7 | 200 | 110 000 |
| CG MARTINI | POPC vesicle (hypertonic) | 60x60x60 | POPC | 4892/3492 | 15.9 | 200 | 120 000 |
| AA | Double bilayer | 10x10x20 | POPC | 300 lipids per bilayer | – | 40 | 25 000 |

[a] initial configuration
[b] after pore closure

#### Supplementary Methods

**Analysis features of Shocker.** The Shocker package has a few in-built analysis methods, which, if enabled, store useful properties of a vesicle each pumping cycle. In the first place, the inner and the outer membrane area as well as the vesicle volume are calculated. It is verified if the structure is whole or if it the crossed periodic boundaries of a box. In the latter case, the smaller pieces are transferred to their original positions to make the structure whole again before further calculations. Subsequently, a point cloud is generated based on the lipid linker positions, in the case of Martini force field only the GL1 beads are considered. Alternatively, the user can provide the desired atom type for analysis. Subsequently, the Python library pyvista [1] was used to generate a triangulated surface of which the area and volume can be determined. This method allows for accurately calculating the area and volume of even the most irregularly shaped vesicles.

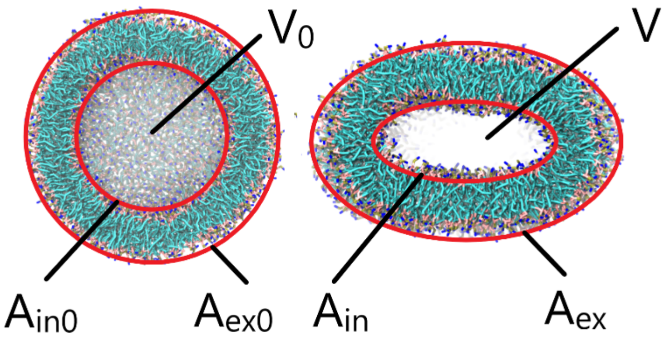


**Supplementary Figure 6 Schematic representation of the different vesicle shape properties analyzed by Shocker.** When a pumping simulation is initialized (left), Shocker determines the initial interior volume ($V_0$), the area of the inner leaflet ($A_{in0}$) as well as the area of the outer leaflet ($A_{ex0}$). At the end of each pumping cycle (right), the current volume (V), the area of the inner leaflet ($A_{in}$), and the area of the outer leaflet ($A_{ex}$) are measured. With these values, Shocker can calculate the relative volume (v) and reduced area difference (Δa) which is used to describe shape changes of vesicles.

In case solutes are present in the simulation box, it is possible to calculate the solute concentration inside and outside the vesicle. For this purpose, the vesicle volume from the previous step and the number of solute molecules are determined. The earlier calculated clusters are used to count the solutes in the inner and outer compartments.

For each pumping cycle, Shocker calculates the area of the inner leaflet ($A_{in}$), the area of the outer leaflet ($A_{ex}$), and the interior volume (V) to calculate the relative volume v and the reduced area difference Δa (see **Supplementary Figure 6**). Hereby, relative volume v is defined as

$$v = \frac{V_0}{V} \quad \text{Eq. 1}$$

and the reduced area difference Δa as

$$\Delta a = \frac{\Delta A}{\Delta A_0} = \frac{A_{ex} - A_{in}}{A_{ex0} - A_{in0}} \quad \text{Eq. 2}$$

where $V_0$, $A_{in0}$ and $A_{ex0}$ denote to the volume, area of the inner and outer leaflet at the beginning of the pumping simulation. A combination of these parameters quantifies the shape of a vesicle of the course of the pumping simulation.[2]

Lastly, Shocker can determine the sphericity parameter Δ and oblicity parameter S. These values indicate the degree of sphericity and oblicity, respectively, of a set of points [3] and are calculated according to the eigenvalues of the gyration tensor

$$T_{\alpha\beta} = \frac{1}{2N^2} \sum_{i,j=1}^{N} (r_{i\alpha} - r_{j\alpha})(r_{i\beta} - r_{j\beta}) \quad \text{Eq. 3}$$

with N being the total number of particles and $r_{i\alpha}$ is the αth component of the position of particle i and α and β = x,y,z. The sphericity parameter Δ and oblicity parameter S are defined as

$$\Delta = \frac{3}{2}\frac{\left[\sum_{i=1}^{3}(\lambda_i - \lambda)^2\right]}{(trT)^2} \qquad \text{Eq. 4}$$

And

$$S = \frac{27\left[\sum_{i=1}^{3}(\lambda_i - \lambda)\right]}{(trT)^3} \qquad \text{Eq. 5}$$

where λ = tr(T)/3. The sphericity parameter Δ is 0 for a perfectly spherical set of points, while S is positive in the case of a prolate shape and negative for an oblate shape.

### Supplementary References


1. Sullivan, C. and A. Kaszynski, *PyVista: 3D plotting and mesh analysis through a streamlined interface for the Visualization Toolkit (VTK).* Journal of Open Source Software, 2019. **4**(37): p. 1450.
2. Käs, J. and E. Sackmann, *Shape transitions and shape stability of giant phospholipid vesicles in pure water induced by area-to-volume changes.* Biophysical Journal, 1991. **60**: p. 825-844.
3. Dima, R.I. and D. Thirumalai, *Asymmetry in the shapes of folded and denatured states of proteins.* The Journal of Physical Chemistry B, 2004. **108**(21): p. 6564-6570.
